# Heparan sulfate expression on B cells modulates IgM expression in aged mice and steady-state plasma cell numbers

**DOI:** 10.1101/514588

**Authors:** Damian L. Trujillo, Nadine Jarousse, Laurent Coscoy

**Affiliations:** Department of Molecular and Cell Biology, University of California Berkeley, Berkeley CA, 94720-3200; Aduro Biotech, Berkeley, CA, 94710; Gilead Sciences, Foster City, CA, 94404

## Abstract

Heparan sulfate (HS) modulates many cellular processes including adhesion, motility, ligand-receptor interaction, and proliferation. We have previously reported that murine B cells strongly upregulate cell surface HS upon exposure to type I interferon, TLR-ligands, or B cell receptor stimulation. To investigate the role of HS on B cells *in vivo*, we utilized EXT1^lox/lox^ CD19-Cre conditional KO mice, which are incapable of synthesizing HS in B cells. We found that suppressing HS expression on B cells has no overt effect in B cell development, localization, or motility. However, we did observe that EXT1 conditional KO mice have decreased poly-reactive IgM in naïve aged mice relative to littermate control mice. Despite this decrease in poly-reactive IgM, EXT1 conditional KO mice mounted a normal B cell response to both model antigens and influenza infection. We also observed decreased plasma cells in EXT1 conditional KO mice after influenza infection. Although EXT1 conditional KO mice have decreased plasma cells, these mice still had comparable numbers of influenza-specific antibody secreting cells to littermate control mice. The findings presented here suggest that HS expression on B cells does not play a major role in B cell development or overall B cell function but instead might be involved in fine-tuning B-cell responses.

## Introduction

Heparan sulfate (HS) is a post-translational modify cation comprised of a repeating disaccharide (uronic acid and N-acetyl glucosamine) that undergoes heterogeneous deacetylation, sulfation, and epimerization [1]. This glycosaminoglycan is attached to a core-protein, creating a heparan sulfate proteoglycan (HSPG) [2]. HSPGs are expressed on the surface of almost all mammalian cells and they modulate many cellular processes including cell adhesion, motility, migration, localization, and ligand-receptor interaction [2–5]. A number of lymphocyte-specific chemokines, cytokines, and growth factors contain HS binding domains (IL2-8, IL10, IL12, IFNγ, TNFα, MIP1β, APRIL, etc.)[5–13], suggesting that HS might be important for the coordination of adaptive immune responses.

B cells are an integral component of the immune system. Through the action of surface and secreted antibodies, B cells are capable of providing both innate and adaptive protection against invading pathogens [14, 15]. This is accomplished largely by the B-1 (CD19^hi^, CD5^+^) and B2 (CD19^intermediate^, CD5^−^) B cell subsets. B-1 B cells secrete poly-reactive IgM in the absence of antigen stimulation ^14, 21^. B-2 B cells are the classic B lymphocytes known to respond to thymus-dependent antigens by undergoing class switch recombination, somatic hypermutation, and clonal expansion. The combined effect of these two populations is that B-1 cell-derived IgM provides general innate-like protection against viral pathogens, while B-2 B cells contribute to the adaptive humoral response and immunological memory. Critical to initiating and maintaining a B-2 B cell response is B cell localization, cell-cell interaction, and responsiveness to numerous soluble ligands (chemokines and cytokines) [16–20]. This is illustrated by the concentration of B cells within the secondary lymphoid organs and their chemokine-dependent migratory patterns that result in the surveying of follicular dendritic cells for antigen [16, 21].

Recently, a number of groups have begun investigating the role of HS in immunity. Using distinct mouse models, Garner *et al.* and Reijmers *et al.* investigated the role of HS expression on B cells [22] [9, 23]. While Garner *et al.* found that HS only modestly affects B cell development, Reijmers and colleagues observed a dramatic phenotype in mice unable to express functional HS on cells of the hematopoietic lineage. In these mice, Reijmers *et al.* observed impairments in B cell development and in the antibody response. A recent report from our laboratory presented evidence that the expression of HS on B cells is tightly regulated [10]. Whereas naïve B cells express very low levels of HS, HS is strongly upregulated on the B cell surface following numerous stimuli that are present during different types of infections (such as IFN-I, toll-like receptor ligands, and B cell receptor-stimulators). Jarousse *et al.* also presented *in vitro* evidence that HS expression on B cells can increase B cell responsiveness to the B cell-specific cytokine APRIL, and as a result, HS-expressing B cells survived longer in culture and produced more IgA [10].

To assess the role of HS expression on B cells *in vivo*, we utilized a B cell-specific deletion of exostosin 1 (EXT1), an enzyme necessary for HS synthesis [24]. In this report, we show that expression of HS is not necessary for B cell development, motility, localization, or homing. We also determined that both the primary and secondary antibody response to infection or antigenic challenge proceeds normally in EXT1 cKO mice. Interestingly however, we show that older EXT1 cKO mice have lower titers of circulating poly-reactive IgM, which is suggestive of impaired B-1 B cell function. We also provide evidence that HS expression on B cells affects either the maintenance or generation of plasma cells induced at steady state, independent of immunological challenge. The data presented here suggest that, although EXT1 cKO mice do not show gross B cell defect,HS expression on B cells might be important for the fine-tuning regulation of antibody secreting cells.

## Materials and Methods

### Mice

All mice were housed and bred in pathogen-free conditions at the American Association of Laboratory Animal Care-approved animal facility at the Life Sciences Addition, or at the Northwest Animal Facility (NAF), both located at the University of California, Berkeley, CA, USA. All animal experiments were approved by the Animal Care and Use Committee at the University of California, Berkeley, CA, USA. C57Bl/6 were obtained from Jackson Laboratory. Actin-CFP and ubiquitin-GFP (C57BL/6-Tg(UBC-GFP)30Scha/J) mice were a kind gift from Professor Ellen Robey. Conditional KO mice were generated by crossing EXT1^flox/flox^ (Jackson labs) mice to CD19-Cre^+^ (Jackson labs) mice (both on the C57Bl/6 background) to produce EXT1 conditional KO (EXT1^flox/flox^ CD19-Cre^+^), and littermate controls (EXT1^flox/+^ CD19-Cre^+^). Breeding occurred using single mating pairs. Mouse pups were weekend 21 days after birth, and caged according to sex. Mice were fed standard pelleted food and water. Experiments were conducted with 6- to 25-week-old mice as indicated, in accordance with institutional guidelines for animal care and use. When Necessary, mice were euthanized by inhalation of carbon dioxide.

### Isolation and culture of B cells

Lymphocytes were isolated from peripheral lymph nodes (inguinal, brachial, cervical, mediastinal, or axillary), the spleen, the bone marrow, or the peritoneal cavity of WT C57Bl/6, littermate control, EXT1 conditional KO, Ubiquitin-GFP, or EXT1 conditional KO-actin-CFP mice. Briefly, a single cell suspension was isolated and treated with red blood cell lysis buffer. For figures 1A, 3, and 4, B cells were purified using the EasySep negative selection mouse B cell enrichment kit (Stem Cell Technologies). B cell purity was assessed by flow cytometry using anti CD19 or B220 antibodies (we routinely observed over 98% purity). Cell viability was verified by 7AAD staining. For culturing of B cells, cells were resuspended at a density of 2×10^6^ cells/ml in RPMI media containing 100mM MEM non-essential amino acids, 55 microM 2-Mercaptoethanol, 1 mM sodium pyruvate and 10 mM Hepes. When indicated, purified B cells were treated with Interferon beta (IFNβ◻◻ at 45U/ml unless indicated otherwise (PBL Interferon source). For evaluation of BM B cell subsets, BM was isolated from mice by flushing the marrow from the femurs. Bone marrow was separated from red blood cells by ficoll prior to staining.

**Figure 1.**
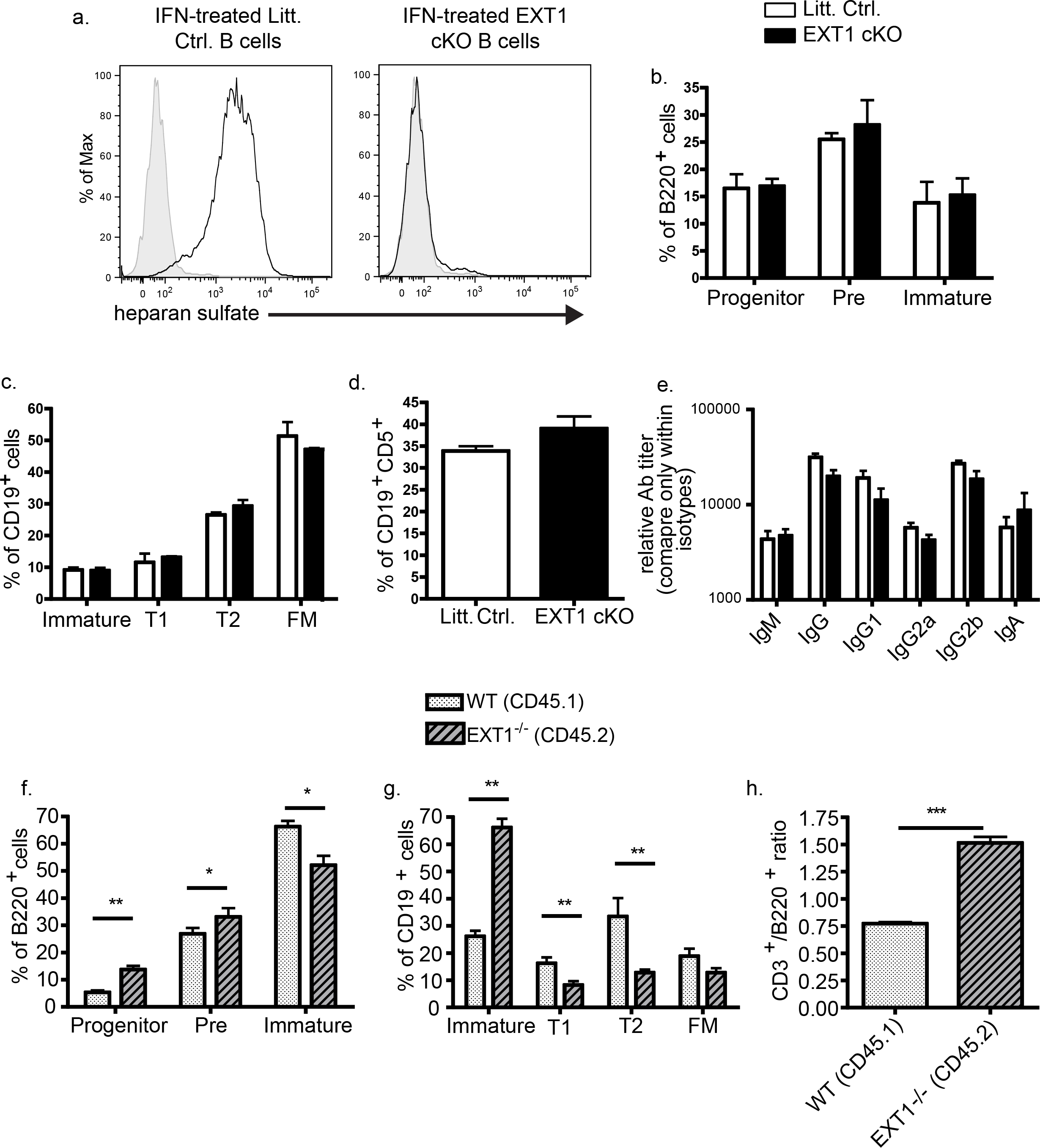
EXT1 expression in B cells plays a subtle role in B cell development. (A) B cells from littermate control or EXT1 cKO mice were purified and cultured for 24 hours in the presence of IFN-I before HS expression was assessed by flow cytometry. Isotype controls are shown as the tinted grey histogram. Bone marrow (B), spleen (C), or peritoneal cells (D) from 6-8-week-old littermate control or EXT1 cKO mice were harvested, stained for B cell populations, and analyzed by flow cytometry. Total antibody serum levels of 6-8-week-old littermate control and EXT1 cKO mice were quantified by ELISA (E). A 1:1 mixture of bone marrow from congenic WT (CD45.1^+^) and EXT1 cKO (CD45.2^+^) were transferred into a lethally irradiated recipient mouse. After allowing 10 weeks for reconstitution, bone marrow (F), spleen (G), or circulating B cells (H) were harvested and stained for B cell populations. Samples were then analyzed by flow cytometry. * or ** indicate *p* values < 0.05 or 0.005, respectively. 1a is one representative experiment from a total of more than 10 repeats. The graphs in 1B-E are experimental averages from 3 experiments where 4-6 mice of the indicated genotype were used per experiment. The graphs in 1F-H are experimental averages from 3 different experiments where 10-30 mice were used per experiment.

### Flow cytometry

Lymphocytes or purified splenic B cells were washed in PBS with 1% BSA and incubated with 1mg/ml purified rat anti-mouse CD16/CD32 (Fc block, BD Pharmingen) at 4°C for 15 min, prior to surface staining. B cells were detected with anti-CD19 or anti-B220 antibodies. HS was detected with the F58-10E4 clone from Seikagaku Corporation. The isotype control was a mouse IgM kappa (TEPC 183, Sigma). Bound antibody was revealed by staining with a FITC- or PE-conjugated anti-mouse IgM^a^ (Igh-6a) monoclonal antibody (BD Pharmingen). Bone marrow progenitor B cells were defined as B220+, CD43 high, IgM low, and IgD low. Bone marrow pre B cells were defined as B220+, CD43 low, IgM/IgD low. Bone marrow immature B cells were defined as B220+, CD43−, IgM/IgD high. Splenic immature B cells were defined as CD19+, IgM−, IgD−. Splenic Transitional type I B cells were defined as CD19+, IgM+, IgD−. Splenic transitional type II B cells were defined as CD19+, IgM+, IgD+. Follicular mantle B cells were defined as CD19+, IgM−, IgD+. B-1 B cells were defined as CD19+, CD5+. Distinction of congenic CD45.1 and EXT1 cKO lymphocytes was done by staining for CD45.1 and CD45.2. For the homing experiments, fluorescently labeled B and T cells were identified with CD19 and CD3 antibodies, respectively. Germinal center B cells were identified as CD19+, Fas+, GL7+, and IgD−. Plasma cells were identified as CD19+, syndecan 1+, and IgD−. For all staining procedures, dead cells were excluded from the analysis by 7AAD staining. Data were acquired on a LSRII flow cytometer (BD Biosciences) and were analyzed with FlowJo software (Treestar).

### Generation of BM chimeras

Wild-type female C57BL/6 congenic mice between 6-8 weeks were lethally irradiated with 900 rads. 24 hours later, the mice were injected intravenously with a 1:1 mixture of congenic WT CD45.1+ and EXT1 conditional KO CD45.2+ (2.5X10^6^ of each genotype) bone marrow. 10 weeks post bone marrow transfer, reconstitution efficiency was checked by tail-vein bleeding and flow cytometry for peripheral B and T cells.

### Homing of B cells to the spleen and peripheral lymph nodes

WT ubiquitin-GFP or EXT1cKO-actin-CFP mice were injected intravenously with either PBS or 0.2 mg Poly I:C. Twenty-four hours later, a single cell suspension of splenocytes were isolated, and red blood cells were lysed. Thirty million splenocytes of each genotype were then injected intravenously into a naïve WT mouse. Twenty minutes or 20 hours later, the indicated secondary lymphoid organs were harvested: at 20 minutes post transfer, spleens were isolated. At 20 hours post transfer, spleen, inguinal-, cervical-, brachial- and axillary-lymph nodes were isolated. A single cell suspension of each organ was isolated, and the ratio of B to T cells from the WT and EXT1 conditional KO donors were assessed by flow cytometry.

### Localization of B cells in secondary lymphoid organs

Littermate control or EXT1cKO mice were injected intravenously with 0.2mg Poly I:C. 24 hours later, a single cell suspension of splenocytes were isolated, and red blood cells were lysed. B cells were purified using the Stem Cell Technologies kit, and labeled with either SNARF (Invitrogen) or CFSE (Invitrogen). 1X10^7^ B cells from each genotype was intravenously injected into a naïve WT mouse. Twenty hours later, the mouse was sacrificed, the inguinal lymph nodes harvested, and cryosectioned. 10μM thick cryosections were transferred onto glass slides and imaged by microscopy.

### Two-Photon imaging of B cells in the inguinal lymph nodes

B cells were purified from naïve littermate control or EXT1conditional KO mice, and labeled with either CFSE or SNARF before 1X10^7^ of each was transferred into a naïve WT mouse. Three hours post transfer, recipient mice were intravenously injected with PBS or 0.2 mg poly I:C. 24 hours post PBS or poly I:C injection, mice were sacrificed, inguinal lymph nodes isolated, and explanted individual lymph nodes were maintained at 37 °C and perfused with oxygenated Dulbecco’s modified Eagle’s medium, and imaged using two-photon microscopy as previously described [25, 26]. Imaging volumes of 172 × 143 × 70–120 μm were scanned every 30 s for 30 min, with 2 μm *z* steps at tissue depths of 30–200 μm below the lymph node capsule using a custom-built two-photon microscope with a Spectra-Physics MaiTai Laser (Newport, Santa Clara, CA, USA) tuned to 920 nm. SNARF, CFSE, and autofluorescence were separated with 495 and 560 nm dichroic mirrors and a 645/75 nm band-pass filter to restrict the detection to the optimal wavelength for each fluorescent protein. For basic motility comparisons between samples, cell tracking was performed using Imaris software (Bitplane, Saint Paul, MN, USA), and speed (path length divided by time) was calculated using MATLAB software (Mathworks, Natick, MA, USA).

### Virus and infections

Purified mouse-adapted influenza A/PR/8/34 (H1N1) was purchased from Advanced Biotechnologies Inc. (Cat. # 10-210-500), aliquoted, and stored at −80 degrees Celsius. For infection of mice, aliquots were diluted in sterile PBS. Mice were anesthetized with isoflurane and infected intranasally with 10^4^ TCID_50_ in a 40ul volume. After intranasal inoculation, mice were allowed to recover on a heating pad. For re-infection/recall experiments, primary infection was performed as described above. Mice were then allowed to recover for a period of 10 weeks before they were re-infected with 5X10^4^ TCID_50_ intranasaly. Mouse survival and weight were monitored daily. As our animal protocol mandates, mice were sacrificed if their weight dropped below 65% their pre-infection weight. For the influenza infection, no mice fell below this percentage of initial weight, though there were some deaths (see Figure 6).

### Titering influenza virus from the lung

At the indicated times, mice were sacrificed and both lungs were harvested in 2 ml PBS. Lungs were then homogenized using a Polytron PT2100 homogenizer (Kinematica). Homogenate was then spun down, and the supernatant was used in an MDCK-agglutination assay as previously described (Cottey, Rowe, and Bender, 2001. (Current Protocols). Briefly, supernatant was serially diluted and added to MDCK cells for 24 hours, and then removed. 5 days later, hemagglutination of chicken red blood cells was used to determine the TCID_50_.

### ELISA for influenza specific antibody isotypes

At the indicated times, mice were sacrificed and a terminal bleed was performed. Serum was separated using serum separator tubes (BD cat.# 02-675-188). ELISA plates were coated overnight at four degrees Celsius with heat-killed influenza (see above), diluted in PBS to 1μg/ml. Serially-diluted serum was added to the plates overnight, before detection of influenza specific antibodies with isotype specific alkaline-phosphotase conjugated antibodies. Substrate conversion (pNpP from Sigma; cat. # N2770) was monitored using a plate reader measuring at λ405nm.

### Statistical Analysis

All statistical results are expressed as mean ± SEM. Statistical analysis was performed using a non-parametric Mann-Whitney test for comparison between two groups (Graphpad software). Differences were considered significant at *p*<0.05.

## Acknowledgements

We would like to thank Professor Ellen Robey, Jenny Ross, and Seong Ji Han, Heather Melichar and Paul Herzmark for help/tutoring with Two-photon imaging and data analysis.

## Conflict of Interest

The authors declare no financial or commercial conflict of interest.

## Results

### EXT1 conditional knock-out mice exhibit normal B cell development

To determine the significance of HS expression on B cells *in vivo*, a conditional knock-out (KO) mouse incapable of expressing HS on B cells was generated. This was done by conditional deletion of EXT1, an enzyme necessary for HS synthesis [24]. Mice with EXT1 flanked by LoxP sites were crossed to CD19-Cre^+^ mice. CD19-Cre^+^ mice express Cre-recombinase under the control of the CD19 promoter, a B cell specific surface protein that is first expressed during the pro-B cell stage of B cell development [27]. Although naïve B cells express little to no HS on their surfaces, HS can be strongly induced by treatment with type I IFN (IFN-I) [10]. As expected, when B cells from littermate control mice (EXT1^fl/+^ CD19-Cre^+^) were treated with IFN-I, high levels of HS expression were induced (Figure 1a, left). Upon IFN-I treatment of B cells from EXT1 conditional KO (EXT1 cKO) mice (EXT1^fl/fl^ CD19-Cre^+^), very little HS expression was detected (Figure 1A, right), indicating that we have generated a mouse unable to express HS on B cells.

Because HS expression has been implicated in lymphocyte development [9],[23], we first determined if the B cell compartment of EXT1 cKO mice developed normally. B cell subsets from mice 6-8 weeks-old within the bone marrow (BM), (progenitor-, pre-, and immature-B cells) were present in comparable proportions to littermate controls (Figure 1B). Similarly, splenic B cell populations (Figure 1C) and B-1 B cells (Figure 1D) from mice 6-8-weeks-old were present in similar proportions to littermate controls. We also examined whether total serum antibody titers were comparable between littermate control and EXT1 cKO mice. By performing ELISA on serum from 6-week-old mice, we found comparable titers of all antibody isotypes assessed in littermate control and EXT1 cKO mice (Figure 1E). The evaluation of the B cell compartment of 6-8-week-old EXT1 cKO mice suggests that HS does not play a major role in B cell development.

### EXT1-deficient B cells exhibit a developmental defect in a competitive context

Although no intrinsic defect in B cell development was observed in EXT1 cKO mice, we seeked to determine if EXT1-deficient B cells developed normally in a competitive context. To address this, we used competitive BM chimeras, which provide a more stringent developmental environment that is capable of revealing small defects. A 1:1 mixture of WT congenic CD45.1^+^ bone marrow and EXT1^fl/fl^ CD19-Cre^+^ CD45.2^+^ BM was transferred into a lethally irradiated WT C57Bl/6 mouse. After allowing 10 weeks for reconstitution, we isolated BM from these chimeras and quantified B cell subpopulations. We observed that relative to congenic WT populations, EXT1-deficient B cells exhibited increased progenitor and pre B cell populations, while having a decreased immature B cell population (Figure 1F). Similar to what was seen in the BM, splenic EXT1-deficient B cells had altered population distributions, with increased immature B cells, and decreased transitional-1 and -2 B cell populations (Figure 1G). The percent of follicular mantle B cells was similar between WT and EXT1-deficient B cells. We also observed an overall defect in B cell reconstitution (Figure 1H, p<0.0001). The above data suggests that HS expression plays a subtle role in B cell development that is revealed only in a competitive context. Because of the observed defects in both reconstitution and B cell development in the mixed BM chimeras, we chose to to look further into B cell behavior and the antibody responses of EXT1 cKO and littermate control mice.

### Older naive EXT1 cKO mice generate lower titers of poly-reactive IgM

As previously shown in figure 1e, 6-week old littermate control and EXT1 cKO mice have comparable titers of serum IgM (Figure 2A). However, naïve EXT1 cKO mice exhibited lower serum IgM titers overtime, relative to littermate controls (Figure 2A). Consistent with a lower serum IgM titer, we detected decreased levels of “polyreactive” IgM specific for the model antigens TNP (Figure 2B), HEL (Figure 2C), as well as influenza (Figure 2D) in aged naïve EXT1 cKO mice. To determine if the decreased IgM observed in older EXT1 cKO mice was a result of fewer B-1 B cells, we quantified peritoneal cavity B-1 B cells. Surprisingly, we found that in older littermate control and EXT1 cKO mice, similar numbers and percentages of B-1 B cells were present in the peritoneal cavity (Figure 2E). This was observed regardless of whether B1B-1 B cells were defined as CD19^+^CD5^+^ or CD19^+^CD5^+^IgM^hi^CD43^+^ (data not shown). These results suggest that the expression of EXT1 in B cells, although unimportant for B cell development, affects the amount of secreted IgM in older naïve EXT1 cKO mice.

**Figure 2.**
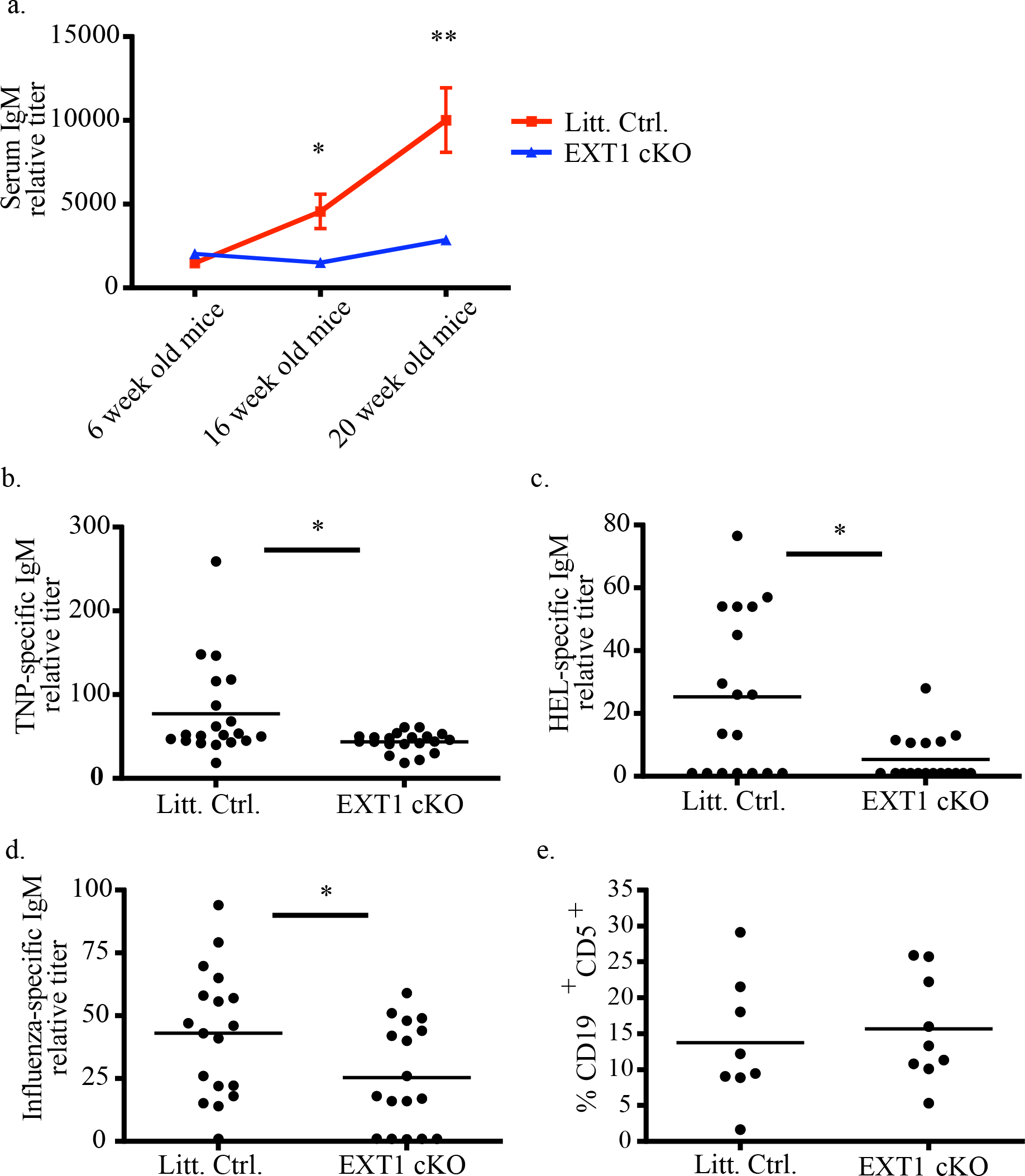
Older naive EXT1 cKO mice generate lower titers of poly-reactive IgM. (A) At the indicated age, serum from EXT1 cKO and littermate control mice was harvested and IgM antibodies quantified by sandwich ELISA. (B-D) Serum from 20-25 week old littermate control and EXT1 cKO mice was harvested and the TNP- (B), HEL- (C), and influenza-specific (D) IgM titers were determined by sandwich ELISA. (E) 20-25-week-old littermate control and EXT1 cKO mice were sacrificed, and the proportions of B-1 B cells (CD19^+^, CD5^+^ double positive) within the peritoneal cavity were determined by flow cytometry. * or ** indicate *p* values < 0.05 or 0.005, respectively. 2A is an experimental average from 4 different experiments where 5-10 mice of the indicated genotype were used per experiment at all three time points. The graphs in 2B-E show the results of 2-3 different experiments where 5-10 mice of the indicated genotype were used per experiment.

### EXT1-deficiency in B cells does not affect B cell homing or localization within peripheral lymph nodes

We next asked if other aspects of B cell biology are dependent on HS expression. HS has been shown to affect cell motility, cell-adhesion, as well as cytokine and chemokine binding [3, 4, 28–30]. Because lymphocyte homing and localization are dependent on the above processes [31], we wondered if B cells lacking HS would home and localize normally. To determine if HS expression affects B cell homing, littermate control and EXT1 cKO mice were injected intravenously (i.v.) with PBS as a control, or poly I:C, which induces expression of HS on littermate control, but not EXT1-deficient B cells [10]. Twenty-four hours later, splenocytes were harvested, differentially labeled with either SNARF or CFSE, mixed 1:1, and transferred into a naïve WT mouse by i.v. injection. Twenty minutes or twenty hours post transfer, recipient mice were sacrificed, and secondary lymphoid organs harvested. The number of transferred B cells relative to transferred T cells per organ was determined by flow cytometry. At both 20 minutes and 20 hours post transfer, we observed that similar ratios of B:T cells from littermate control and EXT1 cKO mice were homing to peripheral lymph nodes and the spleen of naïve mice (Figure 3A-E).

**Figure 3.**
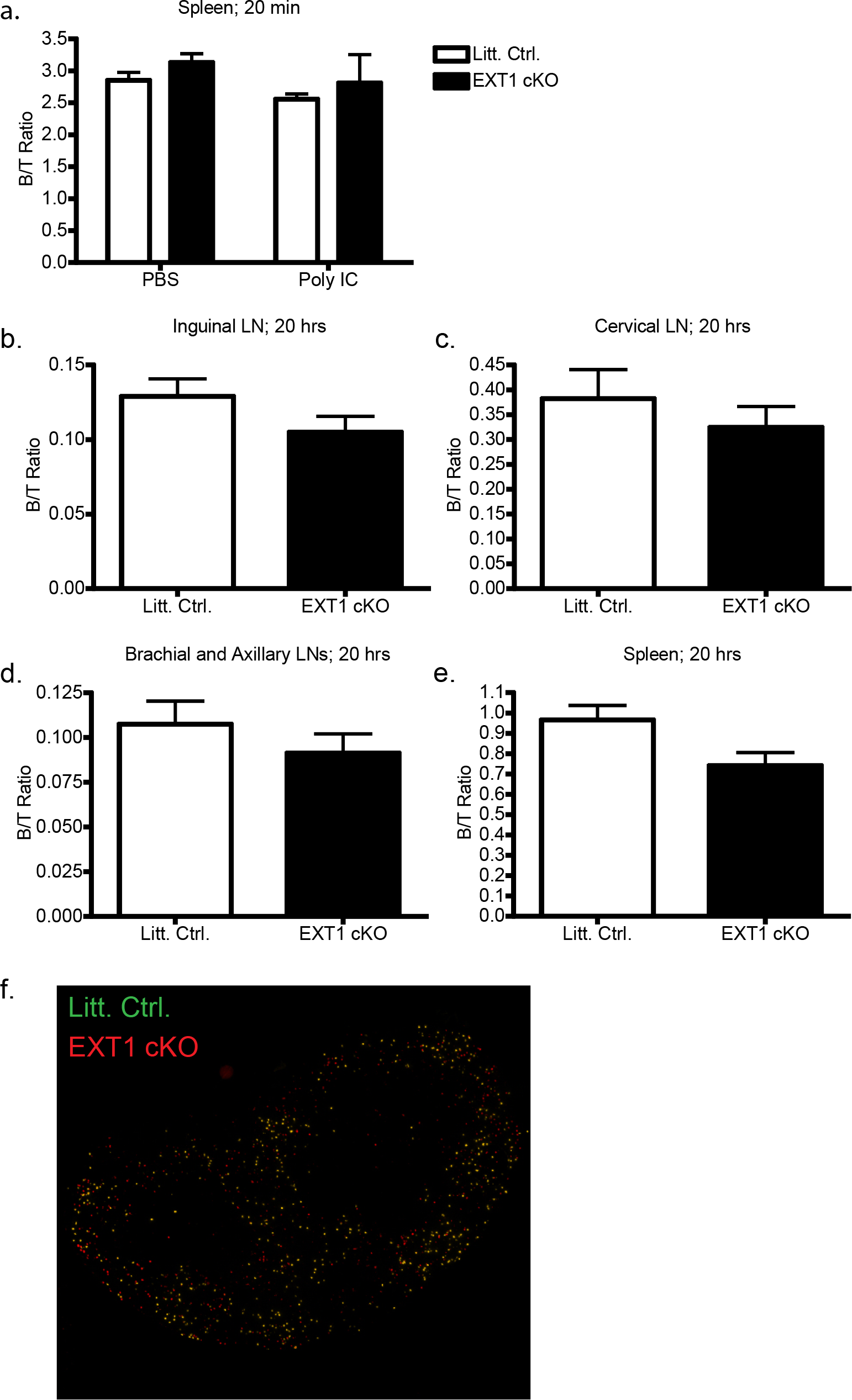
EXT1-deficiency in B cells does not affect B cell homing or localization within peripheral lymph nodes. Splenocytes from PBS or poly I:C treated WT-ubiquitin-GFP mice or EXT1 cKO actin-CFP were harvested and transferred into naïve WT mice. Twenty minutes or twenty hours post transfer, spleens (A and E) and peripheral lymph nodes (B-D) were isolated, and stained for CD3 and CD19. Graphs depict the ratio of transferred B cell:T cell for the indicated organ. (F) Purified B cells from poly I:C treated littermate control or EXT1 conditional KO mice were differentially labeled (littermate control with CFSE, EXT1-deficient B cells with SNARF), and transferred into a naïve recipient mouse. Twenty hours post transfer, inguinal lymph nodes were harvested and cryosectioned and imaged by microscopy. The graphs in 3a-3e depict the averages from 3 experiments where 5 mice received the labeled cells of different genotypes per experiment. Figure 3f shows a representative experiment performed three times with 2-3 mice per experiment.

Using a similar experimental design, purified B cells from poly I:C-treated littermate control or EXT1 cKO mice were labeled and transferred into a naïve WT mouse. Twenty-four hours post transfer, we assessed B cell localization within cryo-sections of peripheral lymph nodes by microscopy. We observed that both CFSE-labeled littermate control and SNARF-labeled EXT1-deficient B cells were distributed evenly throughout the B cell follicle (Figure 3F). The above data suggest that at the times monitored, HS expression on B cells does not affect homing, nor does it affect B cell localization within peripheral lymph nodes.

### B cell motility is unaffected by the absence of EXT1

Because B cell motility within the secondary lymphoid organs is critical for the development of a humoral response [17, 32, 33], we were interested to see if the lack of HS expression on B cells affects motility. To address this, purified B cells from littermate control or EXT1 cKO mice were fluorescently labeled and transferred into a naïve WT mouse, which was subsequently injected i.v. with PBS as a control, or poly I:C (to induce HS expression on littermate control B cells). Twenty-four hours after PBS or poly I:C treatment, inguinal lymph nodes were explanted and imaged by two-photon microscopy (Figure 4A and Video 1). In the presence or absence of poly I:C, we did not observe any difference in B cell motility between littermate control and EXT1-deficient B cells as measured by average speed (Figure 4B-C), directional index (Figure 4D), path length, or cell turning angles (data not shown). Thus, in the conditions monitored, we found no evidence that HS expression on B cells affects motility *in vivo*, neither in PBS-, nor in poly I:C-treated mice.

**Figure 4.**
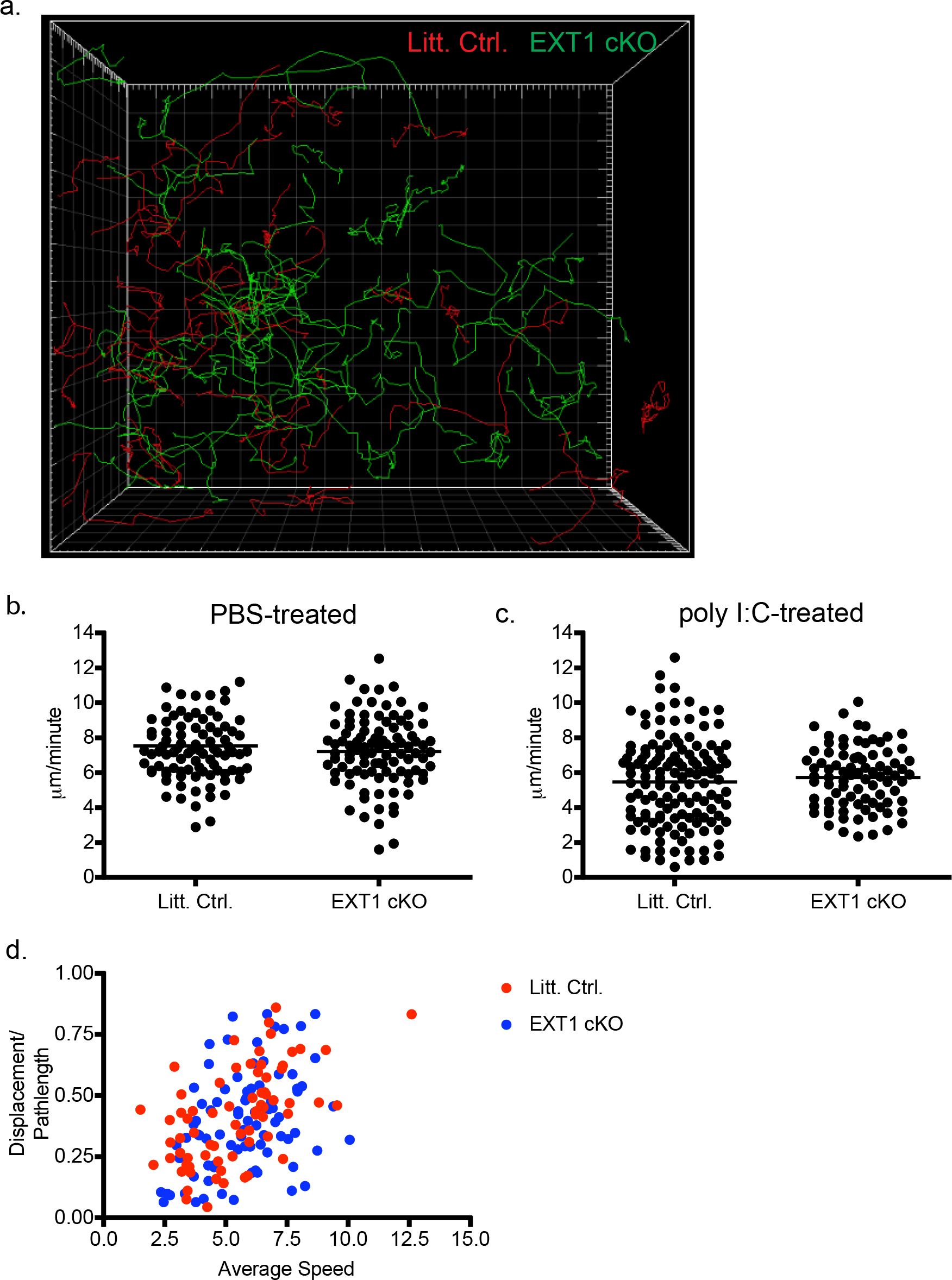
B cell motility is unaffected by the absence of EXT1. B cells from littermate control or EXT1 cKO mice were purified and differentially labeled with SNARF or CFSE. A 1:1 mixture of these cells were transferred into a naïve mouse that was subsequently treated with either PBS as a control, or poly I:C. 24 hours later, inguinal lymph nodes were explanted and imaged by two-photon microscopy. (A) Individual tracks of B cells, either littermate control (SNARF-red) or EXT1-deficient (CFSE-green), from an explanted LN of a poly I:C-treated mouse are shown. The average speed of littermate control and EXT1-deficient B cells was determined for recipient mice treated with PBS (B) or poly I:C (C). Directional path length (D) of individual B cells within a LN from a poly I:C treated mouse, either littermate control (red) or EXT1-deficient (blue), are also shown. Figure 4A shows a representative experiment performed 3 times with 2 mice per experiment. Figures 4B-D show the results of 3 different experiments (with 1-2 mice per experiment) where at least 40 cell tracks where analyzed per experiment.

### Induction of HS expression on B cells after influenza infection

We next decided to assess the importance of HS expression on B cells during a humoral response. To determine if HS expression on B cells plays a role in the humoral response, we chose to use mouse-adapted influenza as an infection model. Influenza was chosen because it is well established that the B cell response is critical for viral clearance and mouse survival [34–36].

Before determining the effects of HS expression on the B cell response, we first assessed HS expression on B cells after infection of 10-week-old mice with influenza. Whereas B cells from uninfected mice express little to no HS (Figure 5A), HS was strongly upregulated on B cells in the lung-draining mediastinal lymph node 48 hours after intranasal infection with influenza (Figure 5B). HS upergulation was transient: HS expression on B cells in the mediastinal lymph node was only slightly above background 14 days post infection (Figure 5C). Interestingly, while germinal center B cells expressed low levels of HS at 14 days pi (Figure 5D), plasma cells expressed a significantly higher amount of HS (Figure 5E). The above data demonstrate that HS is rapidly upregulated on B cells following influenza infection. Furthermore, while HS expression wanes for most B cells post infection, it is still moderately expressed on plasma cells 14 days post infection.

**Figure 5.**
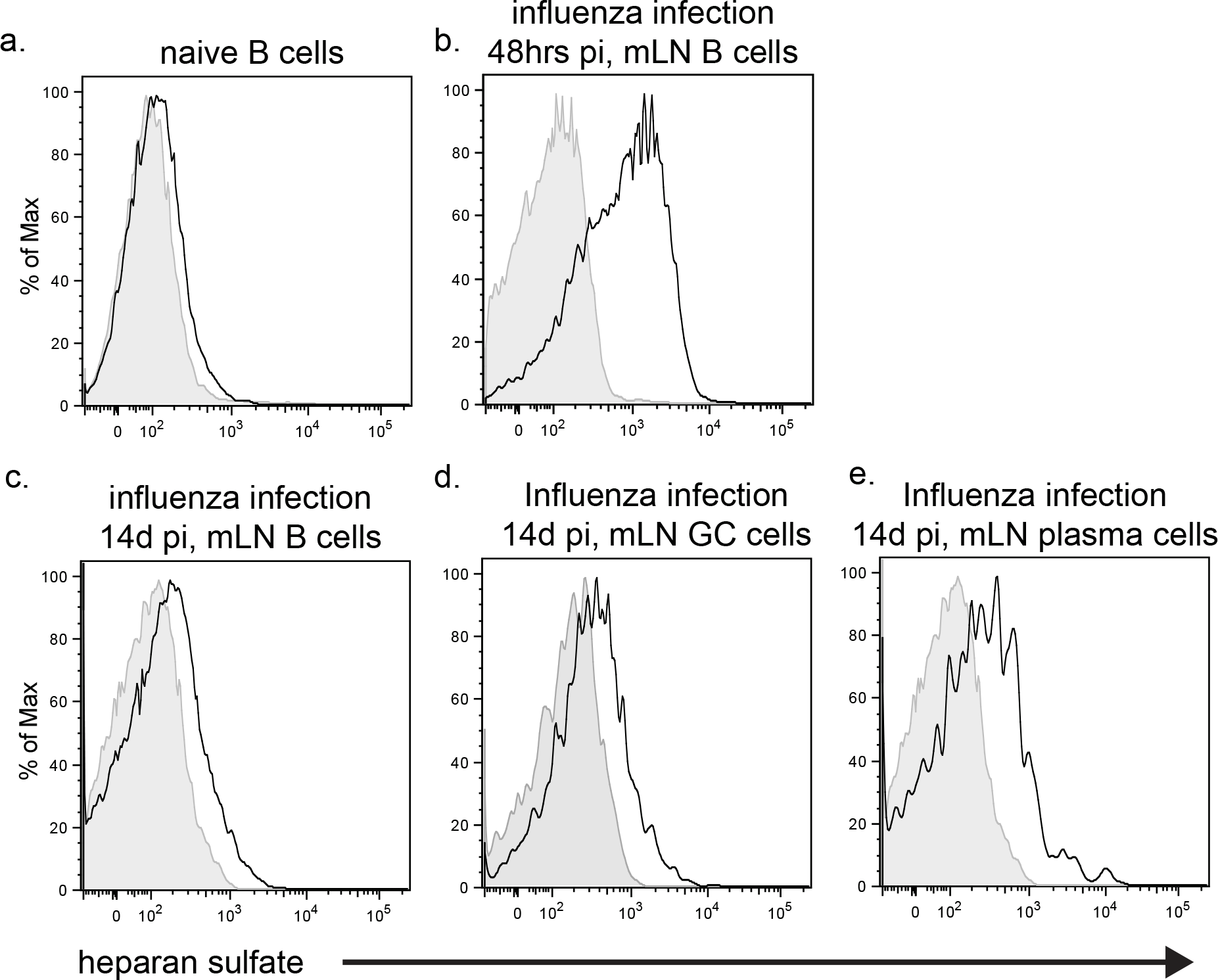
HS is induced on B cells following influenza infection. 10-week old WT C57Bl/6 mice were either left untreated (A), or infected intranasally with 1×10^4^ TCID_50_ A/PR/8 Influenza (B). 48 hours post infection, mediastinal lymph nodes were harvested, cells were stained for CD19 and HS, and HS expression on B cells was assessed by flow cytometry. 10-week-old WT C57Bl/6 mice were intransally infected with 1×10^4^ TCID_50_ A/PR/8 Influenza, and 14 days later, mediastinal lymph nodes were harvested. A single cell suspension of total B cells (C), germinal center B cells (D), and plasma cells (E) were stained for HS. All histograms are representative experiments, using 3-5 mice, and repeated three times.

### EXT1 cKO and littermate control mice exhibit similar morbidity and mortality upon influenza infection

To investigate if the regulated expression of HS on B cells following influenza infection affects influenza-induced pathology or viral clearance, 10-week-old littermate control and EXT1 cKO mice were infected with influenza virus. Following infection, littermate control and EXT1 cKO mice exhibited comparable morbidity, as measured by weight loss (Figure 6A). Similarly, survival rates following infection with 10^4^ TCID_50_ were comparable between the 2 animal groups (Figure 6B). Viral clearance from the lung was also comparable between EXT1 cKO and control mice. (Figure 6C). Thus, lack of HS expression on B cells following influenza virus infection does not affect severity of pathology, nor does it affect the rate of viral clearance from the lung. Similar results were obtained using 25-week old EXT1 cKO mice (data not shown). Thus, despite reports that B-1 B cell-derived IgM is important for early control of influenza [15], we found that even with lower titers of poly-reactive IgM, aged EXT1 cKO mice can still control viral replication in the lung upon influenza infection.

**Figure 6.**
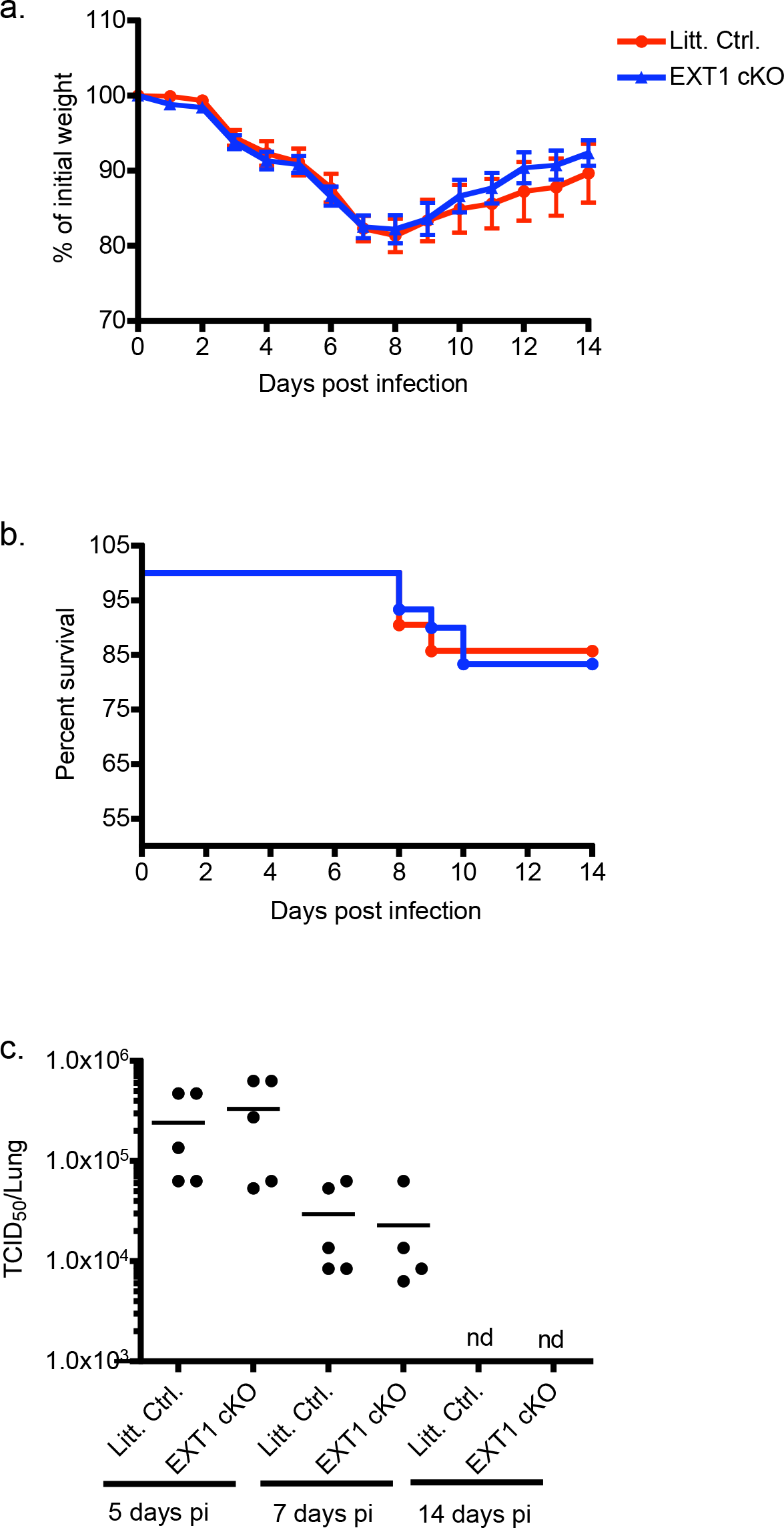
EXT1 cKO and littermate control mice exhibit similar morbidity and mortality upon infection. 10-week-old littermate control or EXT1 cKO mice were intranasally infected with 1×10^4^ TCID_50_ A/PR/8 Influenza. Morbidity as a function of weight loss (A), as well as mouse survival (B), was monitored for two weeks (A). Mice were sacrificed at 5, 7, and 14 days post infection, and the titer of influenza per lung was determined using a MDCK agglutination titering assay. Figures 6A-B depict the averages from 4 experiments where 7-12 mice of the indicated genotype was infected per experiment. Figure 6c is a representative experiment using 5 mice per genotype, and was repeated 3 times.

### EXT1 cKO mice exhibit a decrease in plasma cells after influenza virus infection

Despite seeing no difference in illness severity or viral load when comparing littermate controls and EXT1 cKO mice (aged either 10- or 25-weeks), it is possible that responding B cell populations are affected by the absence of HS expression. We investigated this possibility by harvesting mediastinal lymph nodes and the spleen 2 week post infection of 10-week-old mice. This was followed by enumeration of germinal center (GC) B cells and plasma cells by flow cytometry. While the percentage of GC B cells present in either the mLN or the spleen were comparable between the 2 animal groups (Figure 7A-B), we observed a statistically significant decrease in the percentage of plasma cells in both the mLN and the spleen of EXT1 cKO mice 14 days post infection (Figure 7C-D). The above data suggest that HS expression on B cells plays a role in the generation and/or maintenance of plasma cells.

**Figure 7.**
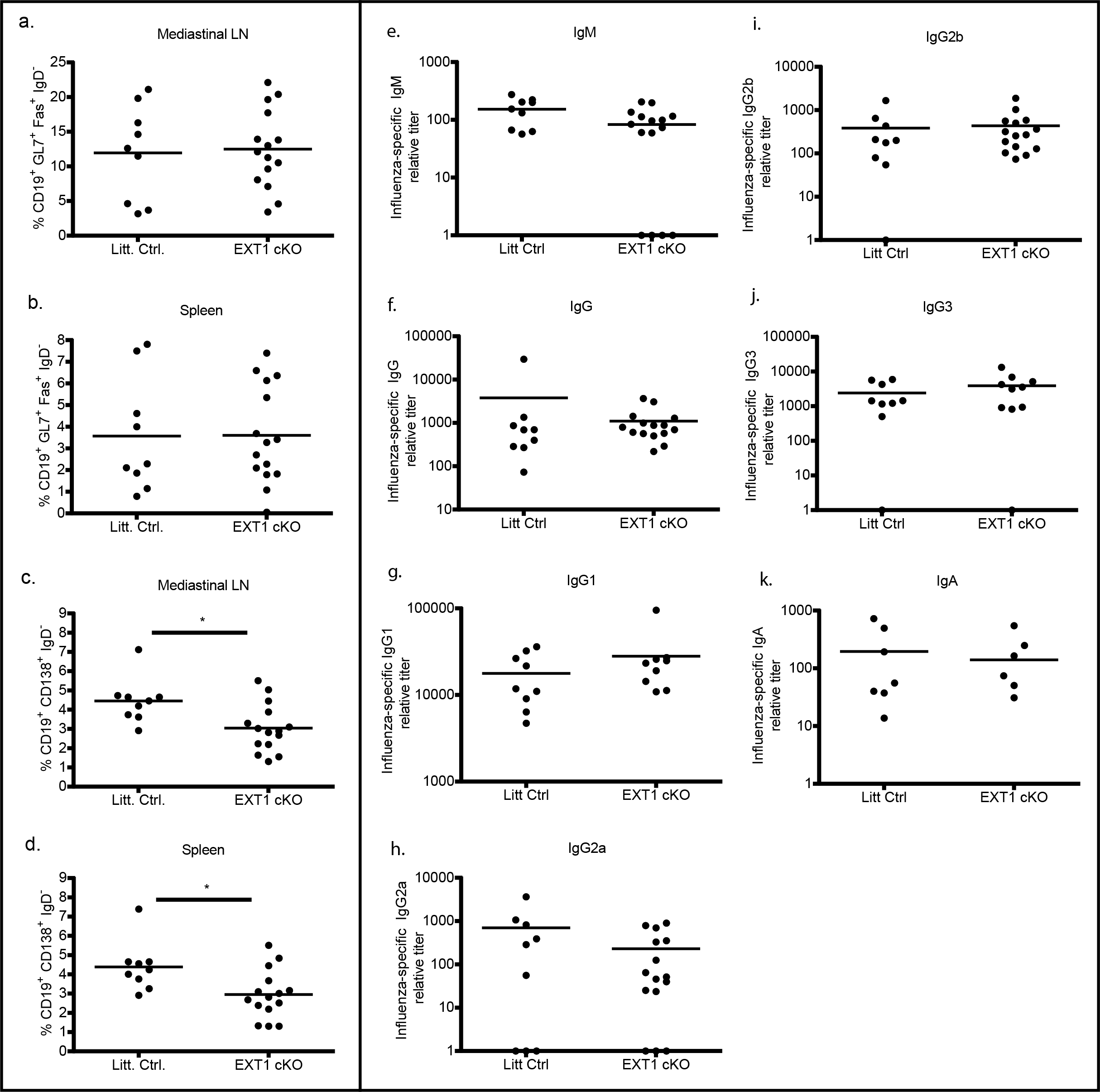
EXT1 cKO mice exhibit a defect in the number of plasma cells generated after influenza infection, however, they generate comparable titers of influenza-specific antibodies. 14 days after infection of 10-week-old EXT1 cKO or littermate control mice with 1×10^4^ TCID_50_ A/PR/8 Influenza, mediastinal lymph nodes and spleens were harvested. A single cell suspension from both tissues was stained for either germinal centers (A and B) or plasma cells (C and D). 14 days after infection of 10-week-old EXT1 cKO or littermate control mice with 1×10^4^ TCID_50_ A/PR/8 Influenza, serum or bronchiolar lavage (BAL) fluid was harvested. Serum (E-J) or BAL (k) fluid was then used in a sandwich ELISA to determine the quantity of different isotypes of influenza-specific antibodies. * indicates p values <0.05. Figure 7A-K show the results of 2-3 different experiments where 3-10 mice of the indicated genotype were used per experiment.

### EXT1 cKO and littermate control mice generate similar titers of influenza-specific antibodies

Having observed a decrease in the number of plasma cells present in EXT1 cKO mice, we seeked to determine if this translated to a reduction in influenza-specific antibodies. To address this, influenza-specific antibody titers in the serum were quantified by ELISA 14 days post infection. For all isotypes evaluated, no differences were observed in influenza-specific antibody titers between littermate control and EXT1 cKO mice (Figure 7E-J). Of particular interest, influenza-specific IgM titers 14 days post infection were comparable between 10-week-old littermate control and EXT1 cKO mice. Because mucosal IgA has also been shown to be important for influenza restriction, we evaluated influenza-specific IgA within the lung. Similarly, we observed no difference between littermate control and EXT1 cKO mice (Figure 7K).

To determine if the decreased plasma cell level in EXT1 cKO mice was unique to influenza infection, 10-week-old (data not shown) and 20-week-old littermate control and EXT1 cKO mice were immunized with the Thymus-dependent antigen TNP-keyhole limpet haemacyanin (KLH). Similar to what we observed upon influenza infection, a lower percentage of plasma cells was detected in EXT1 cKO mice 14 days post-immunization (Supplemental Figure 1B). Despite fewer plasma cells in EXT1 cKO mice, we did not see any reduction in the titers of TNP-specific antibodies in 20-week-old EXT1 cKO mice, either at 14 days post immunization, or 14 days post boost (supplemental figure 1c-e). The only exception was the starting quantity of TNP-specific IgM in aged EXT1 cKO mice (Supplemental figure 1c), which is in agreement with what we observed in Figure 1. These results corroborate what was observed in EXT1 cKO mice infected with influenza virus (Figure 7). The above data suggests that independent of the immunological challenge used (pathogen or model antigen), EXT1 cKO mice are impaired in their ability to either generate or maintain plasma cells. However, this reduction in plasma cells does not translate to lower antigen- or pathogen-specific antibody titers in EXT1 cKO mice.

### EXT1 cKO mice continue to exhibit a reduction in plasma cells after re-infection with Influenza virus

We then examined whether the decreased number of plasma cells in EXT1 cKO mice after primary infection with influenza virus led to a defect in the B cell recall response. Littermate control and EXT1 cKO mice were infected with 10^4^ TCID_50_ influenza. Mice were allowed to recover for a 10 week period before being re-infected with 5X10^4^ TCID_50_ influenza. In agreement with what we observed in Figure 6, EXT1 cKO and littermate control mice exhibited similar morbidity (data not shown) upon primary infection. Upon secondary infection with five times the initial dose, both littermate control and EXT1 cKO mice exhibited reduced morbidity (data not shown), indicative of immunological memory.

Similar to what we observed after primary infection, EXT1 cKO mice exhibited a decrease in the number of plasma cells present in the mediastinal LN and the bone marrow, but not in the spleen (Figure 8A-C). Contrary to what we observed after primary infection with influenza, EXT1 cKO mice also exhibited a decrease in the number of GC B cells in the spleen (Figure 8D). This was unique to the spleen, as there was no statistically significant decrease in GC B cells in the mediastinal LN (Figure 8E).

**Figure 8.**
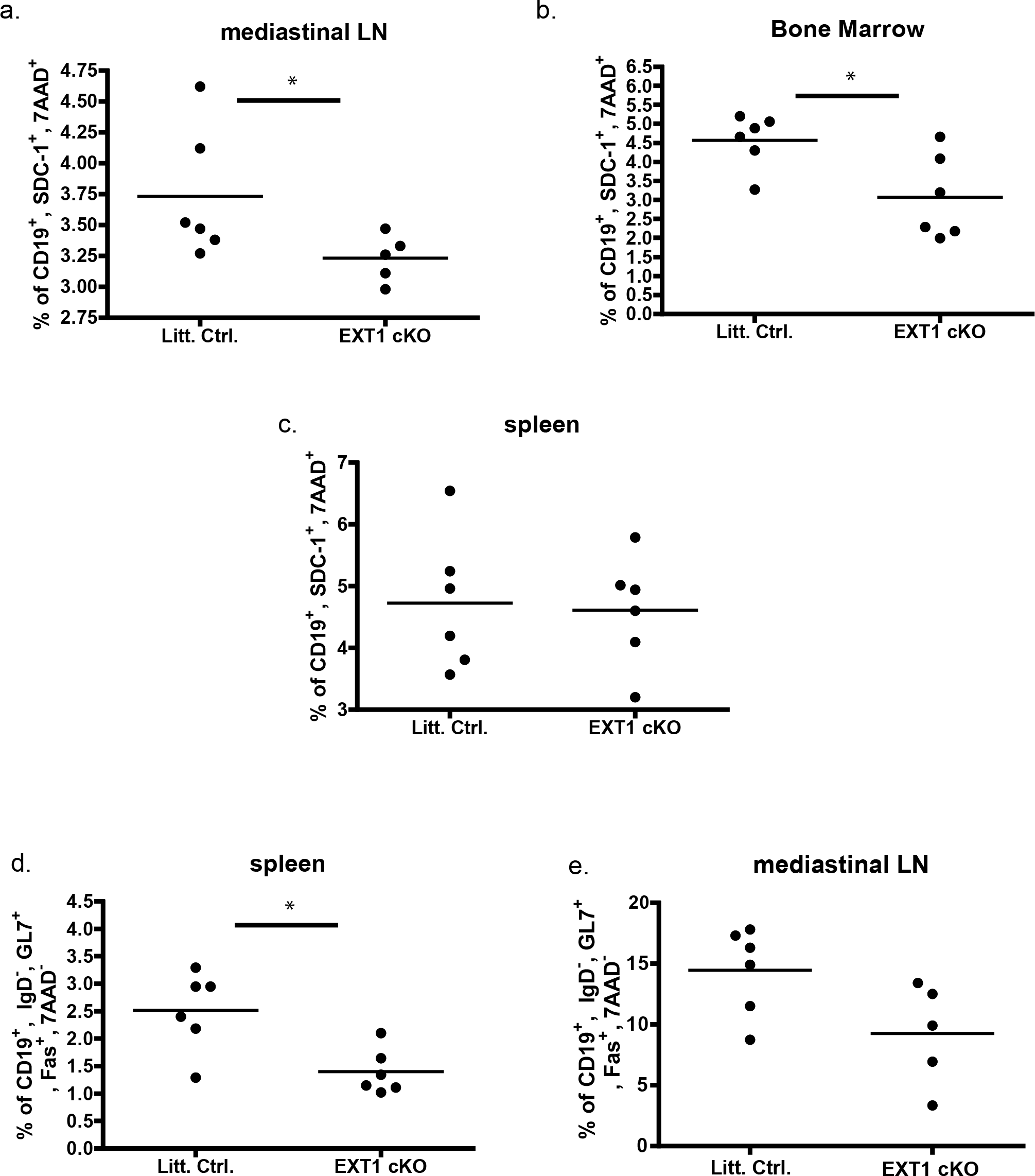
EXT1 cKO mice exhibit a decrease in responding B cell populations after recall infection. Ten weeks after initial infection with a dose of 10^4^ TCID_50_ PR8, mice were re-infected with 5×10^4^ TCID_50_ A/PR/8/34 influenza. 15 days post re-infection, mice were sacrificed, and plasma cells in the mediastinal lymph node (A), bone marrow (B), and spleen (C) were quantified by flow cytometry. Germinal centers in the spleen (D) and mediastinal lymph node (E) were also determined at this time. Figures 8a-8e are representative experiments using 5-7 mice of each genotype, and were repeated 2-3 times.

Influenza-specific antibodies in the serum (isotypes IgM, IgG, IgG1, IgG2a, IgG2b, and IgG3) were quantified at 60 days post initial infection, as well as 7 and 14 days post re-infection. As was seen after primary infection, we detected no difference in the amount of influenza specific antibodies (supplemental figure 2). This was also seen upon recall immunization with the model antigen TNP-KLH (supplemental Figure 1C-E).

Our data indicate that the lack of HS expression on B cells impairs either the generation and/or maintenance of plasma cells after both primary and secondary infections, while not affecting the titers of influenza specific antibodies. To further understand how EXT1 cKO mice (both 10- and 20-week-old mice) have decreased plasma cells, but similar quantities of influenza-specific and TNP-specific antibodies post challenge, we quantified the total number of influenza-specific antibody secreting cells (ASCs). Twenty five-week-old mice were infected with 10^4^ TCID_50_ influenza, and 14 days later, splenocyte and BM cells were harvested to quantify influenza-specific ASCs by ELISPOT. The amount of influenza-specific ASCs was comparable between EXT1 cKO and littermate control mice in both the BM and the spleen (Supplemental Figure 3). The only exception was IgM-secreting influenza-specific ASCs (Supplemental Figure 3G), which is similar to what was seen in Figure 2D. This data suggests that the B cell response generated upon influenza infection, both in antibody titers and in influenza-specific ASCs, is not affected by the lack of HS on B cells. However, the lack of HS does affect the total number of plasma cells, most likely plasma cells that are not influenza-specific.

## Discussion

Numerous studies have shown that HS is important in determining how cells sense and respond to their extracellular environment [3],[37]. We have previously reported that naïve B cells do not express HS, but upon viral or bacterial infection, HS is rapidly upregulated [10]. However, little is known about how HS expression on B cells affects B cell biology. Various groups have presented data suggesting that HS biosynthesis and modification is regulated during B cell development and differentiation, and as a result, HS likely plays a role in B cell biology [38, 39]. To investigate the role of HS expression on B cells *in vivo*, we have characterized a transgenic mouse with a B cell-specific deletion of EXT1, an enzyme necessary for HS synthesis.

### Development and Maturation of B cells in the absence of EXT1

Despite the fact that HS is not expressed from the pro-B cell stage onwards in EXT1 cKO mice, we did not observe any overt defects in B cell development in either the BM or in the spleen. However, a subtle role for HS expression in B cell development was revealed in mixed WT/EXT1 cKO BM chimeras. This developmental defect, observed only in a competitive context, is likely due to the expression of HS slightly above background in BM B cells (data not shown). The alterations in B cell subset populations observed in both the BM and spleen suggest partial blocks at the early stages in B cell development and maturation in the respective organ. In both the BM and the spleen, these partial blocks result in increased precursor B cell populations. These partial blocks suggest that the expression of HS during these stages, although not essential, aids in processes important for B cell maturation. It is tempting to speculate IL-7, a cytokine containing a heparan sulfate binding domain whose signaling is enhanced in the presence of exogenous HS, may account for the defect observed in BM B cell development [6] [40] [41].

### B cell behavior in the absence of EXT1

The homing of B cells to secondary lymphoid organs is key for B cell surveillance of antigens and the development of the humoral immune response [31, 42, 43]. A number of groups have recently demonstrated the importance of HS and sulfated glycan expression on endothelial cells for lymphocyte homing. Both the Fukuda and the Esko groups have shown that the sulfated glycans expressed on endothelial cells serve as ligands for L-selectin expressed on lymphocytes, and interaction between L-selectin and sulfated glycans is important for lymphocyte sticking/rolling along high endothelial venules (HEV). Additionally, presentation of chemokines (such as CCL19 and CCL21) important for integrin-mediated adhesion and transcytosis were diminished in mice deficient for HS on HEVs [29, 44]. Given the importance of HS on HEVs for lymphocyte homing, it was surprising that we saw no defect in B cell homing, either immediate or accumulative, in EXT1-deficient B cells. It remains possible that by assessing only the B:T cell ratio within the secondary lymphoid organs, we may be missing HS-dependent differences in B cell sticking/rolling along the HEVs. Electrostatic repulsion between HS expressed on B cells and sulfated glycans expressed on HEVs may affect B cell rolling. Alternatively, the presence of HS on B cells may alter binding of HEV-presented chemokines such as SDF-1 and CCL19, and as a result, affect integrin activation [29] [31]. Further studies assessing the importance of HS expression on B cells as it relates to B cell rolling and integrin activation may reveal a role for HS in B cell homing.

To the best of our knowledge, we are the first to assess the role of HS expression on B cell localization within secondary lymphoid organs. When looking within peripheral lymph nodes (Fig 3F) and the spleen (data not shown), EXT1-deficient B cells appeared to be evenly distributed throughout the B cell follicle. This was surprising since T cell zone organizing chemokines CCL19 and CCL21 (also known as ELC and SLC, respectively), are known to bind HS [17, 33]. From our data, it does not appear that the presence of HS on B cells enhances responsiveness to these chemokines, as we did not see any difference in the density of B cells at the B cell/T cell border between littermate control and EXT1-deficient B cells. Perhaps only after upregulation of the CCL19/CCL21 receptor on B cells, as occurs upon B cell activation, does HS affect localization within the LN.

Because HS has been shown to affect both chemokine-mediated cell migration [29],[3, 44] and cell motility [45], we were surprised to find no defect in motility in our EXT1-deficient B cells. It is possible that HS expression on B cells can affect motility, however, heparan sulfate or other sulfated glycosaminoglycans present within the extracellular matrix of the lymph node [29] may compensate for the lack of HS on EXT1-deficient B cells. Alternatively, HS expression may affect B cell migration in a context other than that assessed here, or it may only affect B cells within certain parts of lymph nodes such as the border between the B cell follicle and the T cell zone border, where chemokine concentrations are different from those in the B cell follicle [17, 32].

Another facet of HS biology that must be considered is HS composition. That is, both the spacing and the degree to which HS chains are sulfated. A growing body of evidence suggests that sulfation along the HS chain is regulated in a cell- and tissue-specific manner [46],[47]. Importantly, it is now known that distinct sulfation patterns are associated with specific biological activities [37]. Characterization of sulfation patterns along HS chains on B cells may prove key in determining how HS affects B cell biology. Indeed, heparan sulfate modifying enzymes have been shown to be differentially regulated during B cell development and activation [22, 38].

### Aged naive EXT1 cKO mice have lower titers of IgM in the serum

Although we observed no developmental defects in EXT1 cKO mice, we did observe that aged EXT1 cKO mice (over 16 weeks of age) exhibited lower circulating IgM. We were surprised to find that despite lower titers of poly-reactive IgM, EXT1 cKO mice had similar numbers of peritoneal (Figure 2E) and splenic B-1 B cells (data not shown). Although we cannot definitively attribute the lower IgM titers to impaired B-1 B cell antibody production, we hypothesize this is the case based on the poly-reactive nature of the decreased IgM. Furthermore, the B cells that respond to antigenic challenge, B2 B cells, are unimpaired in their ability to respond to both influenza and model antigens (discussed further below). If decreased B-1 B cell antibody production is indeed the cause of the observed lower IgM observed, this would indicate that HS expression on B cells affects function, but not maintenance of B-1 B cells.

How does HS affect antibody production in aged mice? Similar to what was seen in naïve B cells, HS expression on B cells (both CD5-postive and negative) from 6- and 20-week-old mice was slightly above background (Figure 5a and data not shown). Interestingly, Sindhava and colleagues have recently reported that mice lacking APRIL have decreased levels of B-1a and B-1b cells [48]. APRIL has been shown to regulate B cell survival and class switching in a HS-dependent manner [49]. Taken together, this data is in line with our observation that IgM levels are affected in EXT1 cKO mice. We hypothesize that HS and APRIL interact in a way that can regulate B-1 cell numbers and IgM levels. Perhaps the low level of HS expression observed here is key for positioning or responsiveness to factors important for antibody production.

### The B cell response to influenza in the absence of heparan sulfate expression

As was seen with murine gammaherpesvirus 68 (MHV-68) and mouse cytomegalovirus, infection of mice with influenza induces high levels of HS on B cells early post infection [10] (Figure 5B). The strong induction of HS on B cells is likely due to the robust amount of IFN-I that influenza induces in the upper respiratory tract [10],[50]. Also similar to what was observed upon MHV-68 infection, HS expression on most B cells was transient following influenza infection. Notably however, HS expression on plasma cells was still moderately high 14 days pi. The level of HS expression on plasma cells was particularly interesting considering the decreased number of plasma cells present in EXT1 cKO mice post infection (discussed further below).

Both the early thymus-independent, as well as the Thymus-dependent B cell responses contribute to limiting influenza disease severity [36, 51]. Given that HS expression is induced on B cells early after influenza infection, we hypothesized that the early B cell response might be altered, and as a result, this might affect disease severity [15]. We were surprised to find that upon infection with influenza, both littermate control and EXT1 cKO mice tolerated the infection similarly. We also hypothesized that lack of HS expression might affect the generation or maintenance of effector B cells. Indeed, we found that after primary infection with influenza, lack of EXT1 in B cells negatively affects the percentage of plasma cells present. Similarly, we observed a decreased in both plasma cells and GC B cells during the recall response. This observation suggests that while expression of HS on B cells has only a mild affect on GC B cells (only during the recall response), HS expression affects either the generation and/or maintenance of plasma cells. The fact that the decreased percentage of plasma cells did not lead to a decrease in influenza-specific antibody titers is likely due to comparable numbers of influenza-specific antibody secreting cells (Supplemental Figure 3). This suggests that while the immediate generation of influenza-specific antibody secreting cells (ASCs) proceeds normally in EXT1 cKO mice, the total number of plasma cells is reduced.

A possible explanation for this discrepancy is that HS expression on B cell contributes to the maintenance of plasma cells (those plasma cells induced at steady state, independent of influenza infection). As a result, EXT1 cKO mice generate similar numbers of influenza-specific plasma cells, but the non-influenza-specific plasma cells are either not generated or not maintained. In the future, it will be important to delineate the contribution of EXT1 in the generation and or maintenance of antigen/pathogen specific plasma cells over time. The data presented here suggests that the expression of HS on B cells is not necessary to respond to, or clear, the viral pathogen influenza. However, HS expression on B cells does affect the total proportion of effector B cells.

Having observed decreased poly-reactive IgM in aged EXT1 cKO mice, we were surprised that these mice were able to survive and respond to influenza infection comparable to age-matched littermate control mice. given that B-1 B cells secrete both steady state and infection induced poly-reactive IgM. A percent of these IgM molecules function to neutralize influenza during early infection, and they have been shown to be important for surviving an influenza infection in mice^15^. Due to the fact that older EXT1 cKO mice had similar viral titers at 7 and 14 days post infection (data not shown), we believe that even with lower titers of IgM in EXT1 cKO mice, there is still sufficient quantities to control the infection. If we assume that the poly-reactive IgM continues to wane in aging EXT1 cKO mice, we hypothesize that these mice will eventually exhibit defects in controlling influenza. However, one thing to consider is that older mice exhibit impaired immune responses to influenza [52]. Thus, it would be crucial to examine EXT1 cKO mice before they exhibit this age-dependent defect, but when EXT1 cKO are sufficiently old to have minimal poly-reactive IgM.

### Refining our understanding of the importance of heparan sulfate expression on B cells

It is important to note that two other groups have recently investigated the role of HS expression on B cells. Garner and colleagues generated an identical EXT1 cKO mouse, however, they observed a lower efficiency of EXT1 deletion than we report here (Figure 1a) [23]. Similar to the findings presented here, Garner and colleagues also observed no defect in B cell development, with the exception of a slight increase in the number of EXT1 cKO progenitor B cells in the BM. This discrepancy is likely explained by the different genotypes of littermate controls used (EXT1^fl/fl^ CD19Cre^−^ versus our EXT1^fl/+^ CD19Cre^+^). Upon immunization of EXT1 cKO with the model antigens DNP-KLH or DNP-Ficoll, Garner *et al.* observed no defect in the titers of antibodies generated, which is similar to what we observed.

More recently, Reijmers *et al.* investigated the role of GLCE, an epimerization enzyme important for HS maturation [9]. By transferring GLCE-deficient fetal liver hematopoietic stem cells into Rag-2^−/−^γc^−/−^ mice (the authors could not use GLCE-null mice due to embryonic lethality of GLCE-deficiency), Reijmers and colleagues generated mice whose entire hematopoietic compartment is unable to express functional HS. Similar to what we observed in competitive BM chimeras, Reijmers *et al.* reported a defect in B cell reconstitution and B cell development. Because the expression of functional HS on other lymphoid cells was impaired, Reijmers *et al.* could not attribute this defect in B cell development solely to the lack of HS expression on B cells. We saw no developmental defect in EXT1 cKO mouse, and believe what we observed in BM chimeras may be an over estimation of the role of HS in B cell development.

Our observation that lack of HS expression negatively affects the proportion of plasma cells is similar to what Reijmers *et al.* observed after immunization with the model antigen TNP-KLH (Figure 7 and supplemental Figure 1B). Unique to our report is the use of a pathogen, which we show induces high levels of HS expression on B cells. Furthermore, our data allows us to definitively associate the decreased titers of poly-reactive IgM as well as the decreased plasma B cells to EXT1-deficiency in B cells. Given the level of HS expression on plasma cells, as well as the importance of HS expression for responsiveness to the B cell specific cytokine APRIL [10],[9], this result is consistent with the predicted role of HS expression on plasma cells.

Our observation that EXT1-deficiency in B cells does not affect titers of antibodies generated after influenza infection is distinct from Reijmers *et al.*, who found that the amount of antibodies generated after immunization with the model antigen TNP-KLH was decreased. This discrepancy is likely explained by the use of a different mouse model.

### Concluding Remarks

In this report, we provide evidence that the expression of HS plays a nuanced role in B cell biology. While most B cell processes we evaluated were unaffected by the absence of HS, we provide definitive evidence that HS on B cells can affect the quantity of poly-reactive IgM in older mice. Furthermore, we also show that the total proportion of plasma cells is reduced in EXT1 cKO mice. Although this notable reduction in plasma B cells did not impair the ability to cope with the viral pathogen influenza, we hypothesize that HS expression on B cells influences either the generation or maintenance of steady state plasma cells. As a result, we anticipate that it may be key for mounting B cell responses to certain classes of antigens and/or pathogens. Perhaps induction of HS is key in certain types of infections, or perhaps it is important at a certain point in the B cell response that was not assessed here. Although we did not see any defect in the thymus-independent antibody response when EXT1 cKO mice were immunized with NP-Ficoll or NP-LPS (Supplemental Figure 4), HS expression may be key early in the immune response, when HS is most highly expressed on B cells. Alternatively, HS expression on B cells may serve to increase interactions with antigens containing HS-binding domains. Indeed, many viruses are known to encode proteins containing HS-binding domains [53–56]. Further studies examining the response of EXT1 cKO mice to different pathogens, including those that bind HS to gain entry into target cells, may refine our understanding of HS expression on B cells.

