## Supplemental Figure Legend for "Heparan sulfate expression on B cells modulates IgM expression in aged mice and steady-state plasma cell numbers"

**Video 1. Motility of littermate control and EXT1-deficient B cells in the inguinal LN.** A representative movie acquired by 2-photon microscopy of littermate control (labeled in red-SNARF) and EXT1-deficient B cells (labeled in green-CFSE) within the inguinal lymph node of a WT C57B/6 mouse. This movie was the result of a 30-minute time-lapse sequence of 50 μm z-projection images.

**Supplemental Figure 1. EXT1 cKO mice exhibit decreased plasma cells following immunization with TNP-KLH.** 20 week-old Littermate control and EXT1 cKO mice were immunized with 100 μg TNP-KLH in 200 μg poly I.C. intraperitonially. 14 days post immunization, spleens were harvested and germinal center B cells (a), and plasma cells (b) were quantified by flow cytometry. Serum was harvested from littermate control and EXT1 cKO mice prior to immunization, 14 days post immunization, and 14 days post secondary immunization (44 days post initial immunization), and TNP-specific IgM (c), IgG2a (d), and IgG2b (e) antibodies were quantified by ELISA.

**Supplemental Figure 2. EXT1 cKO mice generate comparable titers of influenza specific antibodies pre- and post-secondary infection.** 60 days after primary infection of littermate control and EXT1 cKO mice with $10^4$ $TCID_{50}$, serum was harvested and influenza specific IgM, IgG, IgG1, IgG2a, IgG2b, and IgG3 (a-f) was quantified by ELISA. 70 days post infection, littermate control and EXT1 cKO mice were re-infected with $5X10^4$ TCID50 influenza. 7 (g-l) and 14 (m-r) days post re-infection, serum was harvested and influenza specific IgM, IgG, IgG1, IgG2a, IgG2b, and IgG3 were again quantified by ELISA.

**Supplemental Figure 3. EXT1 cKO and littermate control mice generate similar numbers of influenza-specific antibody secreting cells after influenza infection.** 14 days after infection with $10^4$ $TCID_{50}$ influenza, bone marrow (BM) and spleens were harvested from littermate control and EXT1 cKO mice. Using ELISPOT, influenza specific IgA, IgG and IgM were determined for the bone marrow cells (a, c, e) and the spleen (b, d, f). The total number of antibody secreting cells (IgA-, IgM-, and IgG-secreting) from the bone marrow (g) and spleen (h) are also shown.

**Supplemental Figure 4. EXT1 cKO and littermate control mice generate similar responses to the thymus-independent antigens NP-ficoll and NP-LPS.** Littermate control and EXT1 cKO mice were immunized intraperitoneally with either 20 μg NP-LPS, or with 20 μg NP-Ficoll in conjunction with 200 μg poly I:C. 7 days post-immunization, serum was harvested and np-specific antibody titers were assessed. NP-specific IgM (a and c) and IgG3 (b and d) from NP-LPS- and NP-ficoll-immunized mice were quantified by ELISA, and relative antibody isotype levels are shown.
