## Supplementary figures and images for "Heparan sulfate expression on B cells modulates IgM expression in aged mice and steady-state plasma cell numbers"

### Supplemental Figure 1

#### Supplemental Figure 1

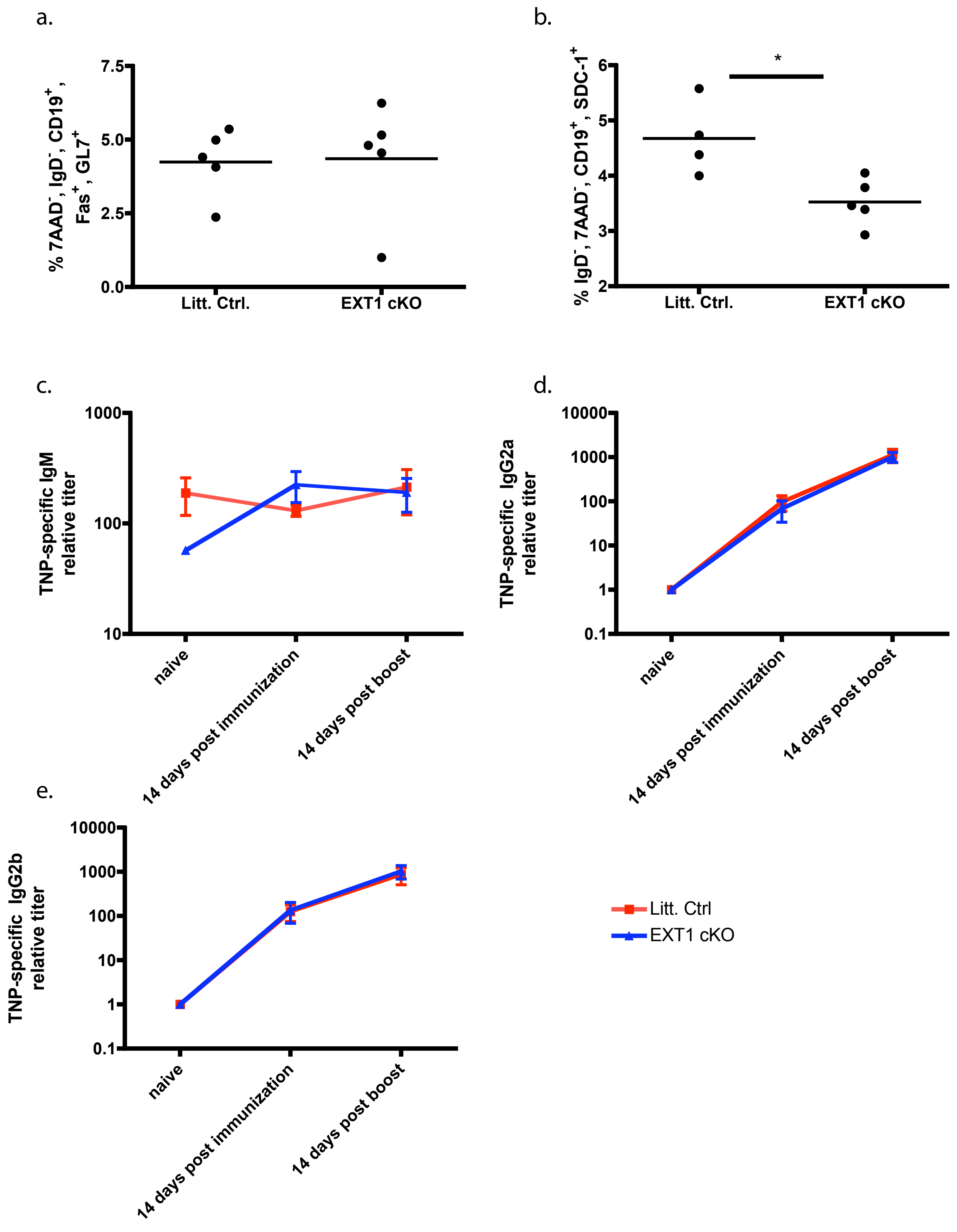

### Supplemental Figure 2

#### Supplemental Figure 2

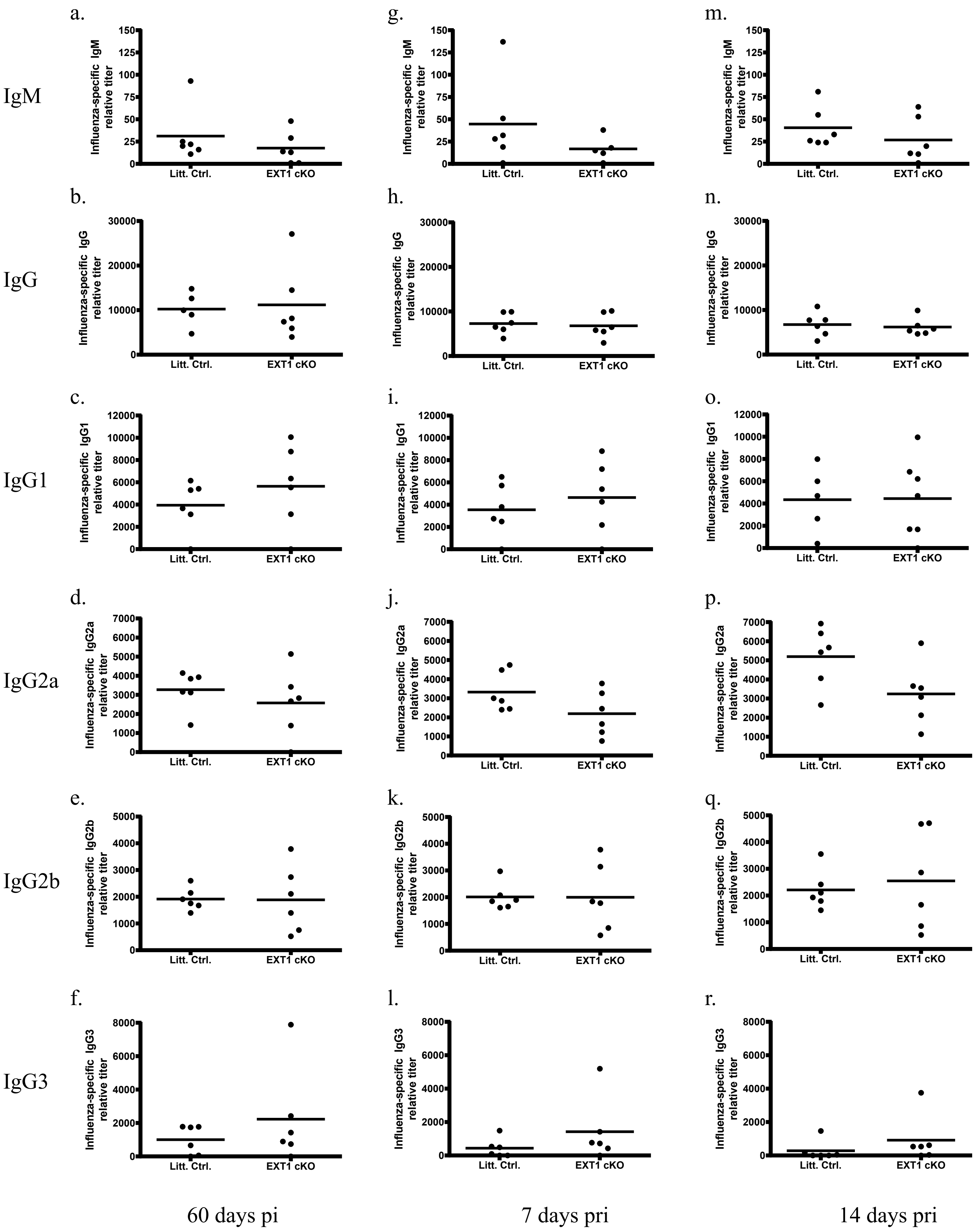

### Supplemental Figure 3

#### Supplemental Figure 3

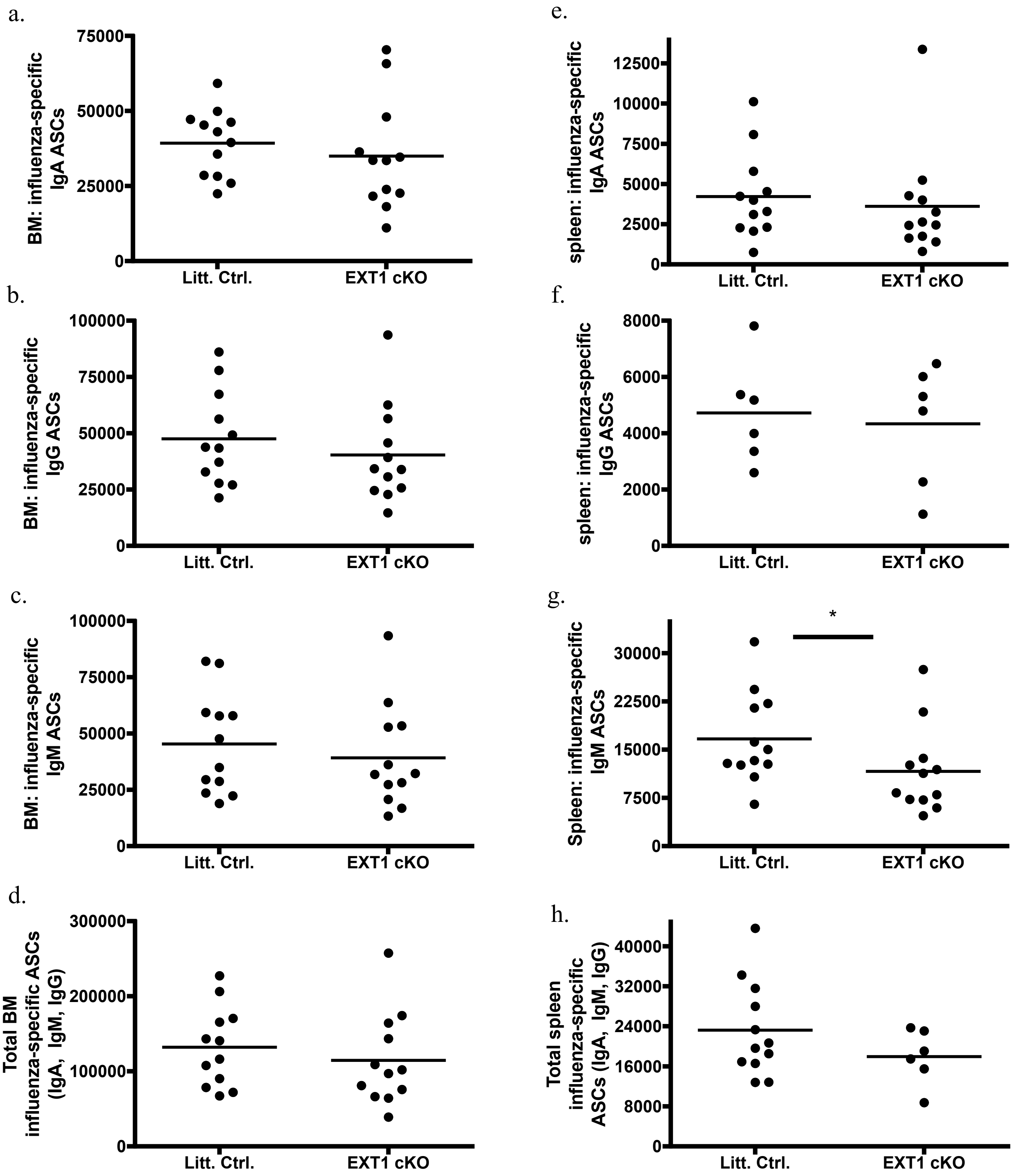

### Supplemental Figure 4

#### Supplemental Figure 4

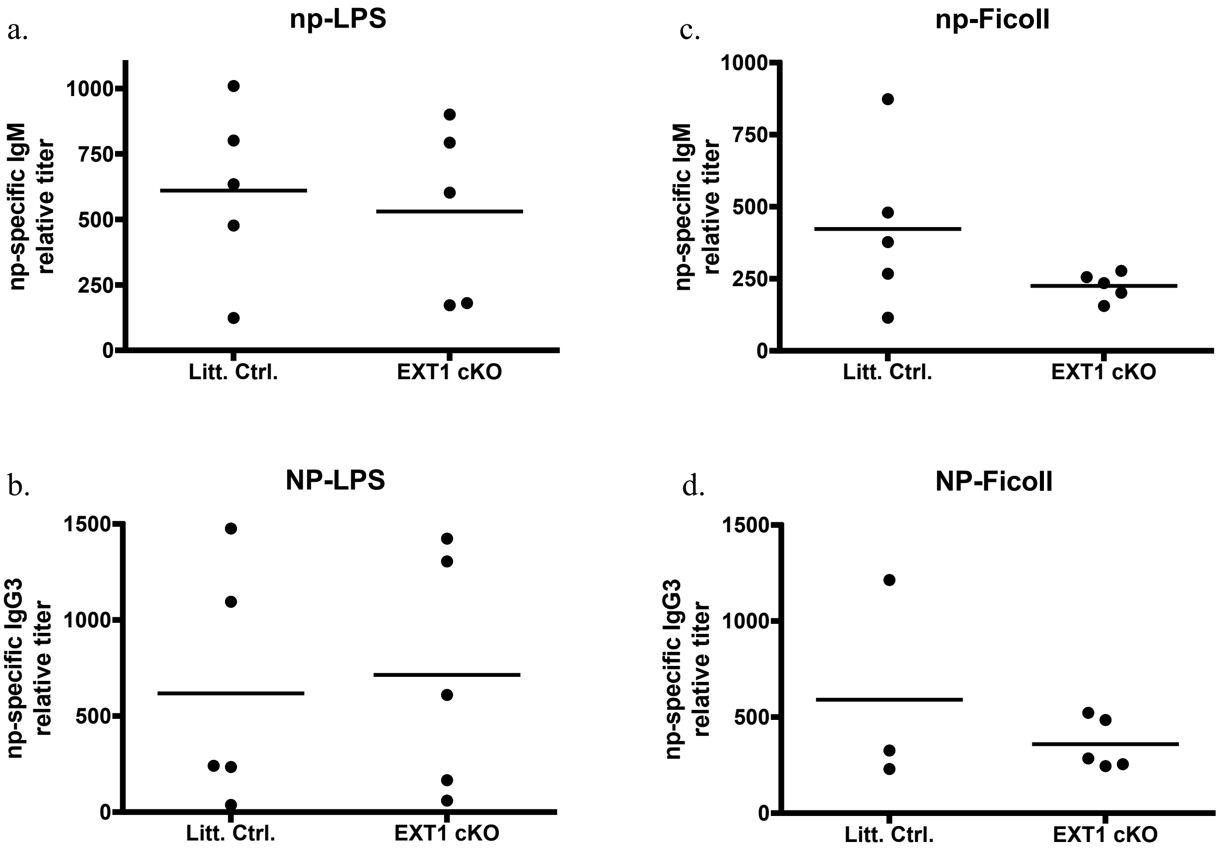
